# Catalytic activity of the prepilin peptidase PilD is required for full *P. aeruginosa* virulence in a nematode infection model

**DOI:** 10.1101/2025.07.10.664072

**Authors:** Jessica M. Cabading, Christopher M. Dade, Katrina T. Forest

**Author notes:** For correspondence: 1550 Linden Drive, Madison WI 53726. These authors contributed equally to this work (Jessica M. Cabading, Christopher M. Dade). Eurofins Microbiology Laboratories, Inc., Madison, WI 53704.

## Abstract

*Pseudomonas aeruginosa* is an ESKAPE pathogen of concern because of its antibiotic resistance and ability to colonize and infect humans in myriad diverse clinical settings, from the lungs of cystic fibrosis patients to burn wounds. Antivirulence strategies have emerged as an alternative to antibiotics for treating *P. aeruginosa* and other pathogens. One proposed antivirulence target is the prepilin peptidase PilD because of its centrality in two virulence mechanisms: the Type IV pili and the Type II Secretion System (T2SS). Substitution of invariant aspartic acids in the putative active site of PilD led to loss of peptidase activity in an *in vitro* cleavage assay and abrogation of both pilus-dependent twitching motility and T2SS-dependent protease secretion. Subsequently, this study utilized a simple *Caenorhabditis elegans* animal infection model to investigate the *in vivo* magnitude of the role of PilD on *P. aeruginosa* virulence. In the absence of functional PilD—either through gene disruption or catalytic inactivation—*P. aeruginosa* exhibited delayed lethality and was reliant on other virulence mechanisms to infect and kill *C. elegans.* These findings highlight PilD as a valuable antivirulence target in *P. aeruginosa*.

**Author summary:** *Pseudomonas aeruginosa* is a tough-to-treat bacterial pathogen that causes serious infections in hospital settings, especially for people with burns, lung disease, or weakened immune systems. As antibiotic resistance grows, researchers seek new ways to stop infections— not by killing bacteria directly, but by blocking the mechanisms they use to cause disease. If a drug could interfere with multiple virulence pathways at the same time, that would make it particularly effective at stopping infection. One possible new drug target to shut down multiple virulence factors is the enzyme PilD, which helps *P. aeruginosa* build two different systems it uses to stick to tissues and secrete harmful proteins. In this study, we tested what happens when PilD is removed or disabled. Using the small, transparent worm *Caenorhabditis elegans* as a simple model for animal infection, we found that without active PilD, *P. aeruginosa* bacteria were much slower to kill their host. Even though the bacteria could still grow, they struggled to attach, spread, and cause damage. These results highlight PilD as a promising target for antivirulence treatments—new types of drugs that disarm harmful bacteria without driving antibiotic resistance. Our findings also support the use of *C. elegans* as a fast, cost-effective system to test potential treatments in living hosts.

## Introduction

*Pseudomonas aeruginosa* is responsible for many hospital-acquired (nosocomial) antibiotic-resistant infections (1,2). It often infects ventilated patients, burn victims, and those with catheters (3). Bacterial colonization of pulmonary tissue is responsible for a high degree of morbidity and mortality in cystic fibrosis, chronic obstructive pulmonary disease, bronchiectasis, and other respiratory diseases worldwide. The best method of treatment is preventing colonization from occurring (4–6). If allowed to persist, *P. aeruginosa* will form chronic and mucoid phenotypes in respiratory infections and typically cannot be eliminated, forcing the patient to undergo decades of antibiotic regimes (7). *P. aeruginosa* is also able to colonize and thrive in burn wounds despite the hostile environment of burn wound exudate (8–10). Therefore, early treatment remains the gold standard to avoid recalcitrant colonization.

Two important virulence pathways utilized by *P. aeruginosa* share a common biophysical mechanism and evolution. During initial host contact, *P. aeruginosa* utilizes the Type 4 Pilus (T4P). This extracellular filament is responsible for twitching motility, adherence to epithelial cells, and dissemination to the blood, liver, kidneys, and lungs (8,11–15). During the acute infection phase, *P. aeruginosa* also secretes dozens of virulence factors, including exotoxin A, LasA and LasB proteases, type IV protease, phospholipase H, and lipolytic enzymes through the Type II Secretion System (T2SS) (16,17). One enzyme, the prepilin peptidase PilD (XcpA), is responsible for processing both the major and minor prepilins comprising T4P and the major and minor endopilins comprising the T2SS (18,19). Prepilin peptidase is an integral membrane (20,21) aspartic acid protease (22). PilD mutants have shown its function is necessary for systemic dissemination and intracellular infection by various pathogens, including *P. aeruginosa* (23,24). Because PilD is essential for two virulence mechanisms that play critical roles in the establishment and acute phases of *P. aeruginosa* infections, it has been proposed as an antivirulence drug target (25). This study aimed to investigate the role of PilD in *P. aeruginosa* virulence using the nematode *Caenorhabditis elegans* infection model, which can mimic a variety of human infection environments, from acute pneumonia to burn wounds (26). We show *C. elegans* can serve as a relatively low cost and high throughput animal model in which to test antivirulence compound leads for *P. aeruginosa* infection.

## Results

### Experimental evidence for Asp149 and Asp213 as aspartic acid protease active site residues

Prepilin peptidase is a multipass integral membrane aspartic acid protease conserved in bacteria and archaea. It is expected that two invariant amino acids at the cytoplasmic side of the inner membrane serve as the catalytic residues (27–29). In some examples, site directed mutagenesis at these positions has abrogated protease function (22,30,31), although such experiments can be complicated when substitutions at other positions also impact PilD function (32), potentially for reasons related to folding, localization or stability. We have previously described production of active prepilin peptidases in a cell free transcription/translation system (28). Here, we created the PilD variant D149A/D213A (PilD_D149,213A_) at the highly conserved Asp sites. To distinguish cleaved and uncleaved pilin in a Western blot band-shift assay, we generated a “long leader peptide PilA” chimera (LlpPilA) by prepending the seventeen amino acid signal peptide from the major pseudopilin GspG (XcpT) from *P. aeruginosa* strain K (PAK) before the mature amino acid sequence of PAK PilA. When co-expressed in the presence of liposomes that mimic the inner membrane of *P. aeruginosa* (33), PilD cleaves the leader peptide from prepilin LlpPilA, while the PilD_D149,213A_ variant cannot (Fig 1A). This result validates D149 and D213 as active site residues in *P. aeruginosa* prepilin peptidase.

**Fig 1.**
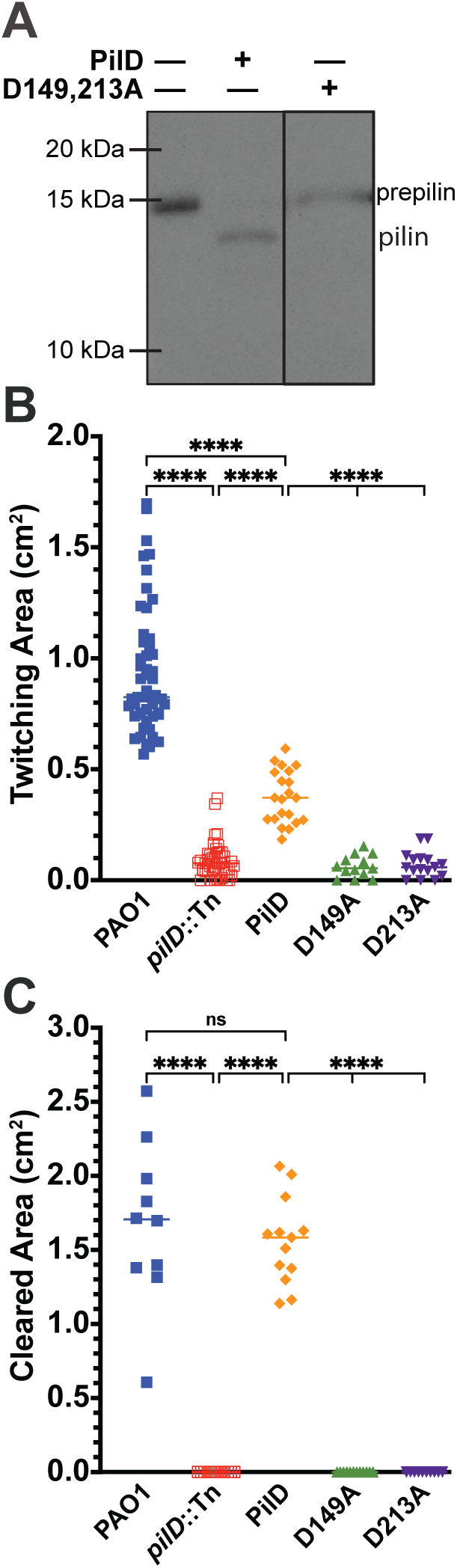
Asp149 and Asp213 are the catalytic residues of PilD and are necessary for twitching motility and T2SS activity. A) When co-expressed in cell-free syntheses, PilD liberates the leader peptide of LlpPilA, while the double substitution PilD_D149,213A_ is unable to. B & C) The inactivation of *pilD* in PAO1 by Tn5 insertion (*pilD*::Tn) or plasmid-based complementation with catalytically inactive PilD variants (D149A or D213A) abrogates *in vivo* catalytic activity. B) T4P-mediated twitching motility and C) T2SS protease secretion comparison. * (p<0.0332), ** (p<0.021), *** (p<0.002), **** (p<0.0001)

#### Catalytically active PilD required for motility and secretion

To investigate the role of *pilD* on *P. aeruginosa* virulence, the effects of catalytic inactivation of *pilD* were investigated via *in vitro* phenotype assays. As has been reported, a transposon insertion into *pilD* of PAK leads to significant decreases in both T4P-dependent twitching motility and T2SS-mediated protease secretion (18,19). This is also true in PAO1 (PAO1*pilD*::Tn) (Fig 1B, C) (see Table 1 for strains and plasmids). Complementation with *pilD* on a constitutive expression plasmid partially restores twitching motility (40% of PAO1 motility) and fully restores T2SS activity (93% of PAO1 activity). In contrast, either substitution D149A or D213A phenotypically copies the PAO1*pilD*::Tn mutant in both assays (Fig 1B, C). A neutral substitution (D151A) had no effect on twitching motility or T2SS activity (Fig S1). Neither catalytic mutant affected growth rate relative to the complemented native *pilD* sequence, although each strain carrying the plasmid had delayed initial growth relative to both PAO1 and PAO1*pilD*::Tn as has been previously reported (34), increasing doubling time from 2.1 h to 3.7 h (Fig S2A, Table S1). Increased doubling time may partially explain why the *pilD* complement does not fully restore twitching motility. These *in vitro* results indicate PilD may be a promising antivirulence target because its absence or inactivation reduces effectiveness of two virulence pathways while not having a marked bacteriostatic or bactericidal effect.

**Table 1.**
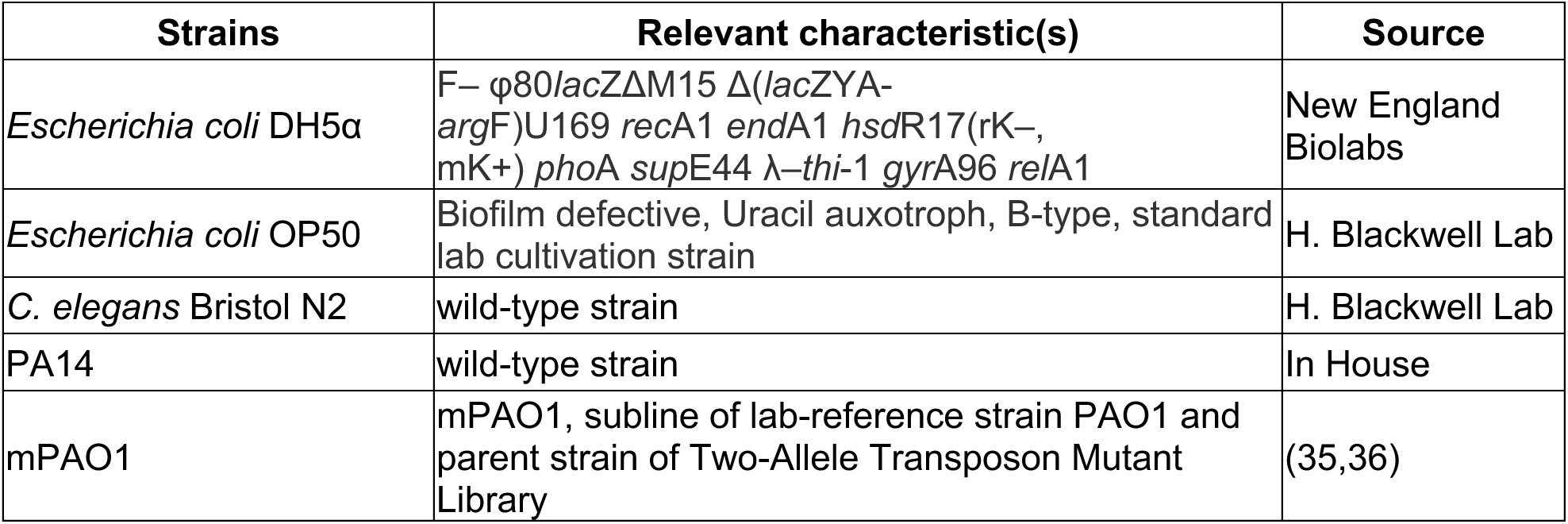
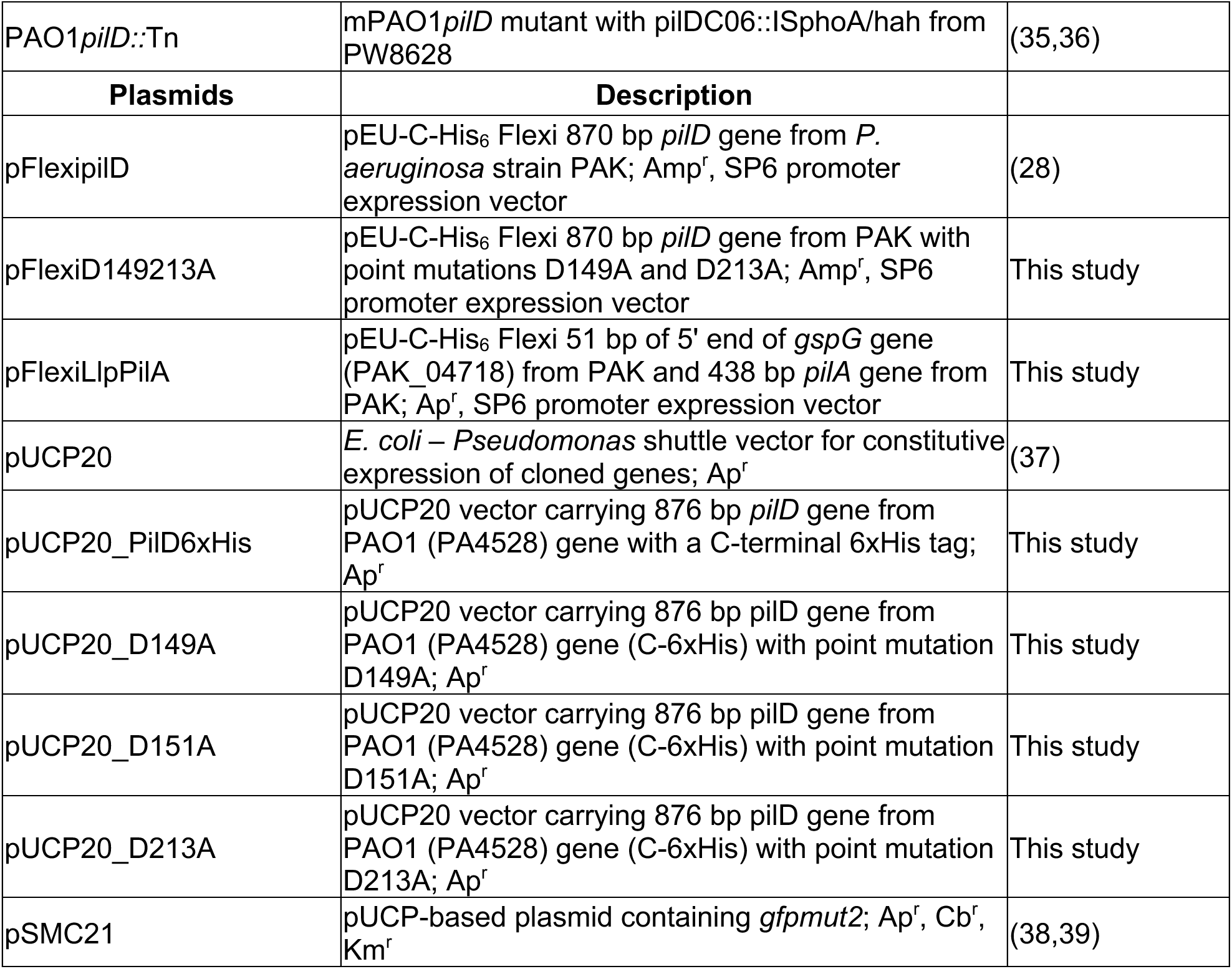

### *pilD* mutant has decreased virulence

In order to establish the simple nematode animal model for *Pseudomonas* virulence (26) in our laboratory, and to assess which *P. aeruginosa* strain would be the best model for PilD-dependent virulence, *C. elegans* nematodes were raised on lawns of PA14 or PAO1, on either high osmolarity peptone-glucose-sorbitol media (PGS) for fast-killing conditions or on low-nutrient nematode growth media (NGM) for slow-killing conditions. *P. aeruginosa* virulence under fast killing conditions is at least partially mediated through low-molecular-weight toxins (40,41), while under slow killing conditions virulence is mediated through gut colonization and proliferation, which is flagella and T4P-depedent (26). As previously observed (26), PA14 killed *C. elegans* extremely quickly under fast killing conditions (supplementary Fig S3A,B, Table S2) potentially due to its unique R-bodies virulence factor that contributes to colonization and pathogenicity in *C. elegans* (42). All subsequent killing assays were performed with PAO1 to isolate the contribution of PilD to *P. aeruginosa* virulence.

To investigate the role of PilD in *P. aeruginosa* pathogenesis under the slow killing regime, nematodes were fed *E. coli* OP50, PAO1 or PAO1*pilD*::Tn grown on NGM. During slow killing, *C. elegans* survival was maintained until approximately 72 hours. After this time point, the *pilD* mutant showed a compromised ability to kill *C. elegans* compared to PAO1 (Fig 2A). The Lt_50_ of *C. elegans* ingesting PAO1 was 120 h. The Lt_50_ increased to 144 h when *C. elegans* ingested the *pilD*::Tn mutant, which was statistically different according to the log rank test (Fig 2C). Because slow killing is mediated through colonization and proliferation, it was expected the *pilD*::Tn mutant would have decreased virulence.

**Fig 2.**
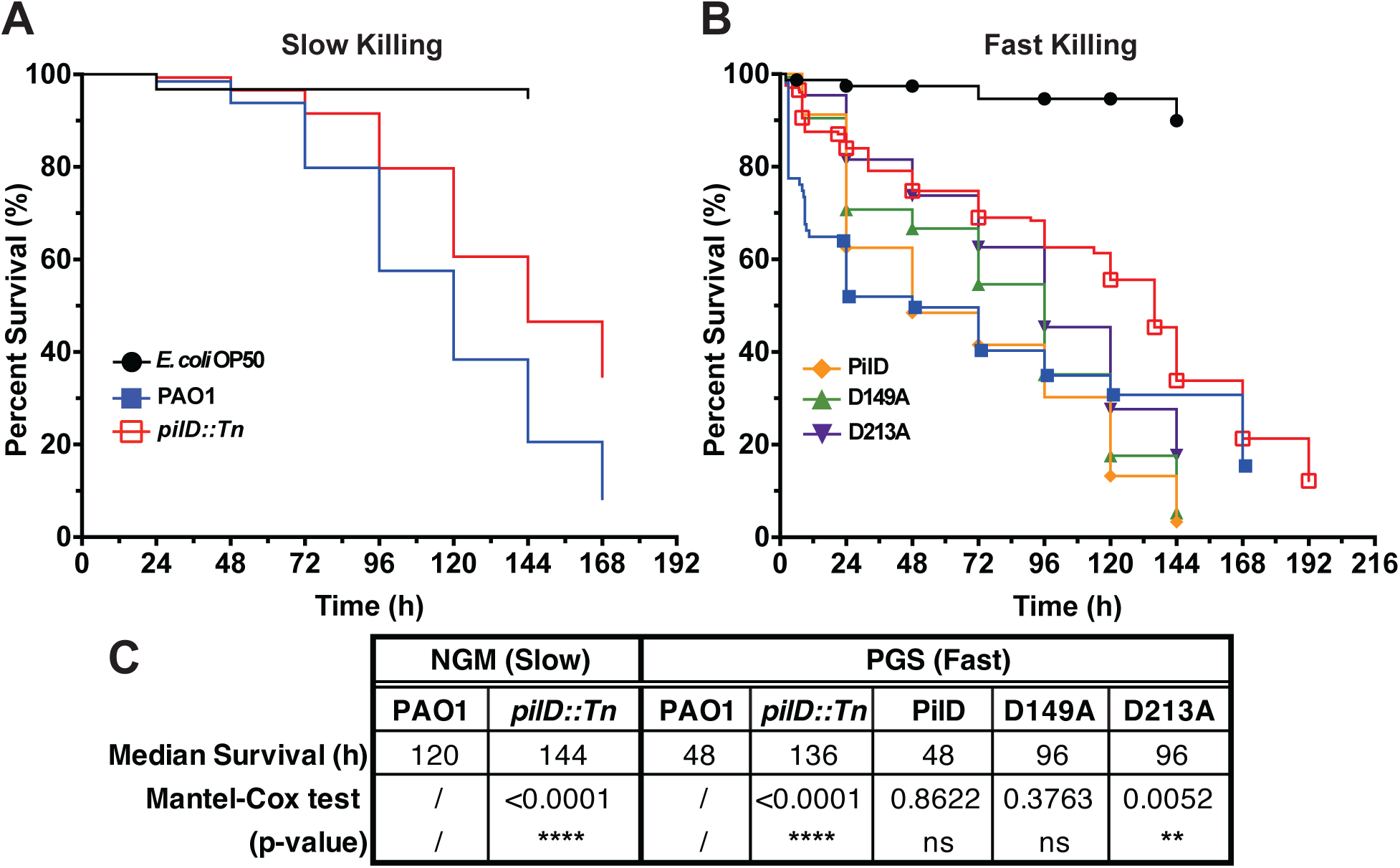
**Prepilin peptidase activity contributes to *P. aeruginosa* virulence during slow and fast killing in *C. elegans***. A) Kaplan-Meier survival curve of *C. elegans* feeding on lawns of *E. coli* OP50, PAO1, or PAO1*pilD::*Tn grown on NGM (slow killing). B) Kaplan-Meier survival curve of *C. elegans* feeding on lawns of *E. coli* OP50, PAO1, PAO1*pilD::*Tn, the *pilD* complemented strain (PilD), and the catalytic aspartic acid point mutant complemented strains (D149A and D213A) grown on PGS. Shapes indicate censored events. C) Median survival (in hours) of *C. elegans* fed on each strain grown on NGM or PGS and the log rank test of statistical significance between kill curves and PAO1 kill curve for each media. * (p<0.0332), ** (p<0.021), *** (p<0.002), **** (p<0.0001)

To explore the potential effect of PilD in toxin-mediated killing under the fast killing regime, nematodes were fed PAO1 or PAO1*pilD*::Tn, and grown on PGS. Within 12 hours, *C. elegans* feeding on PAO1 had significantly lower survival compared to *pilD*::Tn (Fig 2B). The Lt_50_ of *C. elegans* ingesting PAO1 was 48 h, while the Lt_50_ of *C. elegans* ingesting *pilD*::Tn was 136 h (Fig 2C). These killing curves are again significantly different by the log rank test. To verify that change in survival during fast killing was caused by loss of *pilD*, *pilD* was complemented into PAO1*pilD*::Tn on pUCP20 (Fig. 2B). During fast killing, the Lt_50_ of PAO1*pilD*::Tn pUCP20::*pilD* was 48 h, equal to PAO1, and the kill curves were statistically indistinguishable. Complementation with an empty plasmid on the other hand did not restore fast killing (Supplementary Fig S4 and Table S4).

To provide evidence that these effects of *pilD* on *P. aeruginosa* pathogenesis are linked to its proteolytic activity, *C. elegans* was challenged with PAO1*pilD*::Tn strains complemented with *pilD* genes containing the catalytic amino acid substitutions: D149A or D213A (Fig 2B). Both strains had an Lt_50_ of 96 h, which is double the time of PilD and equivalent to the EP Lt_50_ (Fig 2C and Table S4). Compared to PAO1, the D149A kill curve is not statistically significant using the log rank test, though the D213A kill curve is (Fig 2C). Using the alternative Gehan-Breslow-Wilcoxon test, which weights early timepoint data and is preferred for analyzing curves that cross, D149A is also statistically significantly different from (Table S4). These results suggest that, while fast killing is primarily toxin-mediated, PilD activity and thus the T4P and/or T2SS also contributes to *P. aeruginosa* virulence.

#### *pilD* mutant colonizes poorly

To investigate whether the decreased killing efficiency of PAO1*pilD::*Tn was due to a decreased ability to colonize the *C. elegans* gut, nematodes were fed PAO1 or the *pilD::*Tn mutant transformed with the plasmid pSMC21, which constitutively expresses GFP (38,39). Strains were grown on NGM under slow killing conditions to favor gut colonization. Attempts to grow either strain transformed with plasmid pSMC21 on PGS under fast killing conditions were unsuccessful. Corrected total cell fluorescence (CTCF) was determined for each nematode. Compared to PAO1 pSMC21, PAO1 *pilD::*Tn pSMC21 showed lower CTCF (Fig 3A, B). Because the variance within the PAO1 pSMC21 and *pilD::*Tn pSMC21 CTCF measurements were not equal (F-test p-value of <0.0001), Welch’s t-test was used to analyze the statistical significance between gut CTCF measurements, and the difference was not statistically significant.

**Fig 3.**
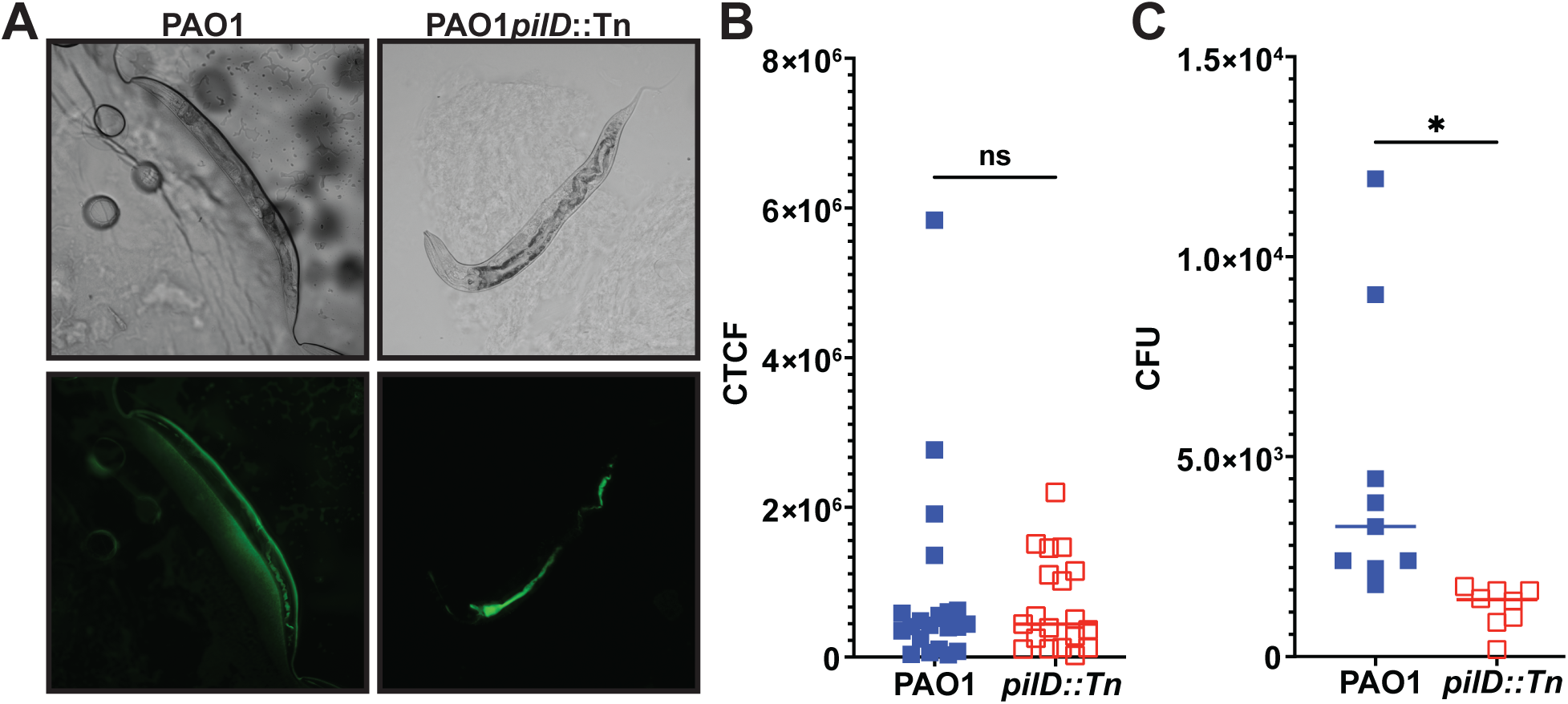
*P. aeruginosa* colonizes the gut of *C. elegans* better than *pilD*::Tn. A) The colonization of the *C. elegans* gut by PAO1 pSMC21 and PAO1*pilD::Tn* pSMC21 can be visualized by fluorescence microscopy. The top panels are bright field images of each representative worm, and the bottom panels are the 488nm channel images of each worm to visualize the GFP signal. B) Corrected Total Cell Fluorescence (CTCF) quantification of gut colonization shows no statistically significant difference in average gut colonization between PAO1–GFP and PAO1*pilD::Tn* pSMC21. C) When the total live-cell bacterial titer of nematode intestines is measured for PAO1 and *pilD::Tn*, the decrease in gut colonization by PAO1*pilD::Tn* is statistically significant (*, p<0.05).

One limitation of whole cell fluorescence labeling is that it cannot distinguish between live and dead cells. Only living *P. aeruginosa* cells, however, can kill *C. elegans* via colonization under slow killing conditions (26). To confirm the colonization trends observed via fluorescence microscopy, we also determined the gut bacterial titer of nematodes fed either PAO1 or *pilD::*Tn. After 24 h, *C. elegans* fed *pilD::*Tn showed decreased colony forming units (CFU) compared to nematodes fed PAO1 (Fig 3C). Because the variances within the PAO1 and *pilD::*Tn CFU measurements were not equal (F-test p-value of <0.0001), Welch’s t-test was used to analyze the statistical significance between average CFU measurements. The difference was statistically significant, confirming *pilD::*Tn is less capable of colonizing *C. elegans* than PA01.

## Conclusion

*P. aeruginosa* is a nosocomial Gram-negative pathogen that threatens the lives of patients with CF, burn wounds, and immunocompromising conditions (43–47). With ever-increasing prevalence of antibiotic resistance, finding alternative treatments is paramount for these patients. In this study, it was shown that PilD plays a role in *in vivo P. aeruginosa* virulence. We also demonstrate *in vitro* and *in vivo* that residues D149 and D213 are the catalytic peptidase residues in PilD and that both are required for twitching motility, T2SS activity (Fig 1), and full *in vivo* virulence (Fig 2).

Two growth media were used to mimic different environments that *P. aeruginosa* might colonize in human hosts. Low nutrient NGM mimics the environment of the lungs during acute pneumonia (48), while high nutrient, high osmolarity PGS mimics the nutrient milieu of burn wounds (9). In both low and rich nutrient conditions, the absence of *pilD* reduces *P. aeruginosa* virulence (Fig. 2). These results extend other *in vivo* studies linking PilD to *P. aeruginosa* virulence in mice (23) and demonstrate PilD-dependent virulence mechanisms contribute to *P. aeruginosa* in diverse infection environments.

As expected under colonization-mediated slow killing conditions, twitching deficient PAO1*pilD::*Tn has decreased virulence. After 72 hours, PAO1 appears to more effectively colonize the nematode gut than *pilD::*Tn, likely because it is able to adhere to the intestinal wall via its functional T4P. The absence of functional T4P in *pilD:*Tn makes it more likely than PAO1 to be excreted by the nematode, thus decreasing *pilD:*Tn colonization and virulence. The presence of functional flagella however, still enables eventual colonization by PAO1*pilD::*Tn and nematode death (8). *P. aeruginosa* additionally contains an arsenal of other virulence mechanisms that contribute to virulence, including LPS, siderophores, and secretions systems I, III, V, and VI (17,49,50).

Somewhat surprisingly, PAO1*pilD::*Tn demonstrated decreased virulence under fast killing conditions as well. Heat-inactivated *P. aeruginosa* cells can still kill nematodes under fast killing conditions (26). Depending on the strain, this virulence has been linked to diffusible small molecule toxins such as phenazines and hydrogen cyanide that are not T2SS-dependent (40,41). In fact, several major T2SS virulence factors, including ExoA, PlcH, and PlcN, have been explicitly eliminated as mediators of fast killing (51). Nevertheless, the decreased virulence of PA01*pilD::*Tn compared to PAO1 manifested within the first 24 hours and persists until 144 hours. The *pilD::*Tn mutant is still able to kill *C. elegans* presumably because it can generate increased levels of hydrogen cyanide and phenazines when grown on PGS. During early-stage infection, however, it is likely that the increased ability of PAO1 to colonize the nematode gut synergizes with toxin production to substantially increase lethality. The decreased early lethality of the *pilD::*Tn mutant even under toxin-mediated killing conditions indicates the value in targeting colonization by *P. aeruginosa* early in infection. PGS media is a more hospitable growth environment than even high nutrient environments, like burn wounds, experienced by *P. aeruginosa* during infection, and the bacteria is likely to rely on a combination of T4P-mediated colonization and toxin production for early-stage virulence during infection.

Mutating either putative catalytic Asp in PilD appears to decrease virulence during fast killing as well, at least during early-stage infection. This indicates PilD peptidase activity specifically contributes to PAO1 virulence *in vivo.* The LT_50_ for both D149A and D213A were twice as long as WT or PilD, although not as long as PAO1*pilD*::Tn.

In the future, it may be illuminating to introduce a genomic *pilD* deletion and catalytic Asp mutations into clinical strains of *P. aeruginosa* to investigate whether lab strain virulence in the *C. elegans* infection model has clinical relevance, especially given the evidence that *xcpA* (*pilD*) truncation mutants are often found in clinical isolates and appear to confer fitness advantages in laboratory competition assays over PAO1 (52). These results indicate that PilD is a promising new antivirulence target, especially for the acute stage of *P. aeruginosa* infection, when paired with wound cleaning or airway clearance techniques to delay or prevent colonization. PilD, however, may be a poor target for chronic infection treatment when the role of T4P in bacterial lifestyle is diminished (53).

## Methods

### Plasmid construction

Plasmids and strains used in this study are described in Table 1. Cultures of *Escherichia coli* DH5α for cloning were grown at 37°C in Luria–Bertani medium (1% peptone, 0.5% yeast extract, 0.5% NaCl). 100 µg/mL carbenicillin was added for plasmid propagation. Point mutations were introduced in pFlexipilD, previously constructed in our lab for cell-free PilD synthesis (28), via sequential two-fragment Gibson assemblies to generate pFlexiD149213A (54). To generate pUCP20 constructs, *pilD* was first amplified by polymerase chain reaction (PCR) from mPAO1 genomic DNA incorporating 20–25 bp overlaps with the adjacent target vector sequences and ligated via a three-fragment Gibson assembly with two pUCP20 vector backbone fragments split to enable scarless reassembly of the ampicillin resistance gene. Single point mutations were introduced via two fragment Gibson assembly as above.

To generate the LlpPilA chimera, *gspG(xcpT)* was first PCR amplified from PAK genomic DNA and ligated via a three-fragment Gibson assembly as above into pET28b+. Then, 51 bps of *gspG* encoding the seventeen amino acid signal sequence were PCR amplified with the 5’ pET28b+ vector backbone. Concomitantly, the 438 bps segment of PAK *pilA* encoding the mature PilA amino acid sequence was PCR amplified from a pET28b+ construct with the 3’ pET28b+ vector backbone incorporating sequence homology with the 5’ 51 bps of *gspG*. These two fragments were ligated together using two-fragment Gibson assembly as above to generate pET28bLlpPilA. To generate pFlexiLlpPilA, the Llp*pilA* chimera gene was first PCR amplified from pET28bLlpPilA, and a three-fragment Gibson assembly was used as above with two pFlexi backbone fragments split within the kanamycin resistance gene, with the 3’ vector backbone fragment lacking the C-terminal His tag.

Plasmids were purified using the Wizard Plus SV Minipreps DNA purification System (Promega), according to the manufacturer’s instructions. Whole plasmid sequencing was performed by Plasmidsaurus using Oxford Nanopore Technology with custom analysis and annotation.

#### Strains

To obtain plasmid-carrying lines, PAO1*pilD*::Tn cells were made chemically competent as previously described (55). These cells were then transformed with either an empty pUCP20 vector or with each *pilD* construct or with pSMC21 (55). Briefly, 100ng of purified plasmid were added to 30µL of chemically competent cells and iced for 1h. Cells were heat shocked at 37°C for 3-5 minutes and immediately iced for 2 minutes. 500 µL of SOC media was added to cells and then incubated at room temperature for 10 minutes. Cells were then incubated at 37°C 250 rpm for 1 h before plating on 250 µg/mL carbenicillin LB agar plates and incubated overnight at 37°C.

#### Cell-free protein synthesis and PilA cleavage band shift assay

Cell-free synthesis of PilD, PilD_D149A,D213A_, and LlpPilA was carried out as previously described (28) with slight modifications. Briefly, transcription and translation reactions were set up using the Wheat Germ Protein Synthesis kit (WEPRO® 7240 Expression Kit). The translation reaction was carried out using the bilayer method. A 25 μL reaction mixture containing 10 μL of WEPRO® 7240, 10 μL of transcription reaction mRNA solution, and 5µL of 50mg/mL liposomes (6:2:2 POPE:POPG:CL) was overlaid with a 200 μL SUB-AMIX® SGC solution in a 96-well polystyrene plate (Greiner), sealed, and incubated at 18°C for 72 h. For co-translations, PilD and LlpPilA transcription mRNA solutions were added in a 1:9 µL ratio. Liposomes were prepared by initially drying a lipid mixture in chloroform (1-palmitoyl-2-oleoyl-sn-glycero-3-phosphoethanolamine (POPE), 1-palmitoyl-2-oleoyl-sn-glycero-3-phospho-(1′-rac-glycerol) [sodium salt] (POPG), *E. coli* cardiolipin [CL], molar ratio 6:2:2) (Avanti Research) by evaporation under a nitrogen stream. The lipidic film was then hydrated with SUB-AMIX® SGC and bath sonicated for 30 min. Liposomes were extruded 11 times through a 0.2μm filter (Avanti Research) and flash frozen in aliquots stored at −80°C until needed for experiments. Whole reactions were then mixed with 6x sample dye buffer before loading on 15% SDS-PAGE (sodium dodecyl sulfate polyacrylamide) gel and visualized via Western blot as previously described with slight modifications (28). Pilin antiserum was used in a 1:5,000 dilution with appropriate secondary antibodies. Western blot imaging was performed with the SuperSignal™ West Pico PLUS Chemiluminescent Substrate (Thermo Scientific) on the Bio-Rad ChemiDoc MP Imaging System using auto optimal imaging settings for chemiluminescent and colorimetric channels to build a composite image.

#### Twitching and T2SS protease secretion assays

Twitching assays were performed as previously described with slight modifications (56). Single colonies were picked with a sterile wooden toothpick and stabbed vertically through the thin layer of LB agar in a twitching assay plate (10mL in a 60mm petri dish), and plates were incubated at 37°C for 24 h. The agar was then removed, and biofilms were stained with Coomassie blue and imaged. The radius of twitching was measured with Fiji (57). Zones were assumed to be circular, and the twitching area was calculated using equation 𝐴 = 𝜋𝑟^!^.

T2SS protease secretion assays were performed as previously described with slight modifications (58). Single colonies of each strain were picked and grown in LB media supplemented with appropriate antibiotics overnight at 37°C 250 rpm. The OD_600_ of each culture was measured and normalized to 0.3 with LB media. 5µL of each culture was spotted onto a 1.5% w/v skim milk LB agar plate and incubated at 37°C for 24 h. Plates were imaged, and the radii to the edge of each colony and cleared zones were measured with Fiji. The area for each was calculated as for the twitching assay assuming a circular area. The area cleared by each strain was then calculated by subtracting the area of the bacterial colony from the area to the edge of the cleared zone, if present.

#### Growth Assays

For growth assays, overnight cultures for each strain were inoculated in LB as above at 37°C 250 rpm. The OD_600_ of each culture was then measured and normalized to 0.1 with LB media. 200µL of each culture were then incubated in a 96-well polystyrene plate (Greiner) at 37°C, and the OD_600_ was measured every 10 min for 6 h in a Tecan M200 pro plate reader with 1mm orbital shaking between reads. Growth wells were surrounded by wells filled with LB media to minimize edge effects and evaporation during incubation and performed in triplicate.

### *C. elegans* Maintenance and Age Synchronization

*C. elegans* were maintained at 18°C on NGM plates seeded with *E. coli* OP50 as previously described (59,60). NGM was made by autoclaving 3g NaCl, 17g agar, and 2.5g peptone in 975mL H_2_O. After the media cooled in a 55°C bath, 1mL 1M CaCl_2_, 1mL 5mg mL cholesterol in ethanol, 1mL 1M MgSO_4_, and 25mL 1M KPO_4_ buffer were added (60). *C. elegans* experimental populations were age synchronized prior to each study following a modified protocol (60). Gravid adults were bleached and eggs transferred to NGM plates seeded with *E. coli* OP50 and incubated at 20°C. At L4 stage, worms were transferred to experimental plates.

#### Killing Assay

For slow and fast killing assays, feeding plates were prepared as previously described (26,41,61). Briefly, PAO1 and PAO1*pilD::*Tn were streaked on LB agar, and PAO1*pilD::*Tn carrying pUCP20 plasmids were streaked on LB agar supplemented with 100μg/μL carbenicillin. Individual colonies were picked and grown in LB media (without antibiotics) overnight at 37°C 250rpm. 150μL of each culture were spread on NGM or PGS plates and grown for 24 hours at 37°C followed by 24 hours at 20°C. PGS media was made with 5g Bacto protease peptone, 5g NaCl, 13.7g sorbitol and 8.5g of agar in 450mL of sterile water. After the media cooled in a 55°C bath, 25mL 1.5M sorbitol and 25mL 20% D-glucose were added.

20-30 age-synchronized nematodes were then transferred to a pathogen lawn plate. Worms were scored as alive or dead by stimulating movement by stroking the worms with a 32-gauge platinum wire. If worms burrowed into the agar, left the bacterial lawn, died of bag of bodies phenotype, or survived to the end of the study, they were scored as censored. Nematodes were transferred to fresh assay plates every 24 hours.

#### Fluorescent Microscopy

All microscopy was performed on the Nikon N-STORM/PALM at the University–Wisconsin Madison Biochemistry Optical Core. PAO1 pSMC21 and PAO1*pilD*::Tn pSMC21 fluorescence were verified by visualizing the bacteria under an epifluorescence microscope. Nematodes were age synchronized as above. L4 stage nematodes were washed and transferred to NGM or PGS plates with lawns of PAO1 pSMC21 or PAO1*pilD*::Tn pSMC21 prepared as above. Nematodes were visualized at 24 h for bacterial gut accumulation. Infected nematodes were transferred using a 32-gauge platinum wire and immobilized in a drop of 1mM levamisole on a 3% agarose pad placed on a microscope slide as described previously (62). Corrected total cell fluorescence (CTCF) within the gut of *C. elegans* was quantified using ImageJ (63).

#### Gut Colonization Colony Forming Units Quantification

Bacterial gut colonization quantification was performed as previously described with slight modifications (64). Nematodes were age synchronized as described above. L4 stage nematodes were washed and transferred to NGM or PGS plates with lawns of PAO1 or PAO1*pilD::*Tn. After 24 h, 20 nematodes were transferred using a 32-gauge platinum wire to 500mL of 1mM levamisole to paralyze the nematodes and inhibit defecation of bacteria. Nematodes were then superficially sterilized with 1% commercial bleach for 1 minute and then centrifuged for 1 minute at 1,300 x *g*. 300mL of supernatant were removed and replaced by 300mL of 1mM levamisole. The solution was centrifuged again at 1,300 x *g*. This was step was repeated 3 times. Nematodes were then homogenized for 1 minute to liberate bacteria colonizing the gut, and the solutions were diluted with sterile water to 500 µL. Samples were serial diluted 1/10, and 50µL of each dilution was plated on Pseudomonas Isolation Agar plates. Plates were incubated for 24 h at 37°C, and plates with 30–350 colonies per plate were analyzed to calculate colony forming units.

#### Statistical Analyses

Prism (version 10.4.2) was utilized for all statistical analyses and graphing. For twitching and T2SS protease secretion assays, one-way ANOVA was used to compare the means of either twitching areas or cleared zones, respectively.

Survival outcomes were plotted on Kaplan-Meier curves, a nonparametric survival estimator which determines the probability of survival over a specific time range while taking into consideration difficulties associated with *in vivo* studies such as censoring (65). Curves were analyzed with the log rank test (Mantel-Cox test) to compare survival distributions for *C. elegans* grown on each experimental strain against PAO1. Because the survival curves of several strains crossed the WT curve and the log rank test is unlikely to detect survival curve differences (66), the Gehan-Breslow-Wilcoxon test was also performed, which gives greater weight to earlier time points (67) (Table S4).

For fluorescence microscopy CTCF and gut titer CFU measurements, the means of each strain were compared using Welch’s t-test because it was found that their variance was not equivalent (F-test p<0.0001).

## Acknowledgements and Funding

We thank Dr. Helen Blackwell at UW–Madison for providing the B2 *C. elegans* animals, Dr. Betty Slinger at UW–Madison for *C. elegans* growth and infection guidance and Dr. George O’Toole at Dartmouth College for providing the pSMC21 plasmid. We acknowledge funding from the UW– Madison Foundation via the E.B. Fred Professorship to KTF and the Ira L. Baldwin Fellowship to CMD. This project was also supported in part by a fellowship award to CMD through the National Defense Science and Engineering Graduate (NDSEG) Fellowship Program, sponsored by the Air Force Research Laboratory (AFRL), the Office of Naval Research (ONR) and the Army Research Office (ARO) and through the UW-Madison Foundation and an Advanced Opportunity Fellowship awarded to JMC through the SciMed Graduate Research Scholars at University of Wisconsin– Madison.

## Author Contributions

KTF was responsible for project conceptualization and funding acquisition. JMC performed experimental investigations and analyses related to *in vivo* pathogenesis. CMD carried out all *in vitro* experiments. All authors contributed to data interpretation and visualization. JMC drafted manuscript; CMD and KTF contributed to writing, editing and finalization of manuscript.

